# Intergenic regions of Saccharomycotina yeasts are enriched in potential to form transmembrane domains

**DOI:** 10.1101/2022.10.21.511897

**Authors:** Emilios Tassios, Christoforos Nikolaou, Nikolaos Vakirlis

## Abstract

Intergenic genomic regions have essential regulatory and structural roles that impose constraints on their sequences. But regions that do not currently encode proteins, also carry the potential to do so in the future. De novo gene emergence, the evolution of novel genes out of previously non-coding sequences has now been established as a potent force for genomic novelty. Recently, it was shown that intergenic regions in the genome of *S. cerevisiae* harbor pervasive cryptic potential to, if theoretically translated, form transmembrane domains (TM domains) more frequently than expected by chance, a property that we refer to as TM-forming enrichment. The source and biological relevance of this property is unknown. Here we expand the investigation into the TM-forming potential of intergenic regions to the entire Saccharomycotina budding yeast subphylum, in an effort to explain this property and understand its importance. We find pervasive but variable enrichment in TM-forming potential across the subphylum, regardless of the composition and average size of intergenic regions. This cryptic property is evenly spread across the genome, cannot be explained by the hydrophobic content of the sequence, and does not appear to localize to regions containing regulatory motifs. This TM-forming enrichment specifically, and not the actual TM-forming potential, is associated, across genomes, with more TM domains in evolutionarily young genes. Our findings shed light on this newly discovered feature of yeast genomes and constitute a first step towards understanding its evolutionary importance.

## Introduction

Genomic regions between genes, known as intergenic regions or intergenes, contain promoters, enhancers and other regulatory sequences and can also have roles related to chromosomal structure. Apart from any roles they may presently have however, they also carry a theoretical potential: to acquire protein-coding capacity in the future through a process now known as de novo gene emergence. Under this evolutionary phenomenon, an entirely novel gene or an entirely novel domain of a preexisting gene emerges out of a previously non-coding region^1,2^. De novo gene emergence has now been recognized as an important and widespread force for evolutionary novelty^3,4^.

By definition, a de novo emerged gene will inherit properties and features that existed in the ancestral non-coding sequence. It follows that, characteristics of intergenic regions, or at least some of them, may govern the initial characteristics of de novo emerged novel genes, and possibly influence their evolutionary trajectory. Multiple studies have demonstrated that young genes (with proven de novo origin or otherwise) have properties that either reflect those of intergenic regions or place them at an intermediate stage between intergenes and conserved genes^5–11^. Such findings substantiate a wider evolutionary link between intergenic regions and young genes. Yet, other studies have described cases where young genes tend to have exaggerated characteristics of conserved genes rather than intergenes, as for example has been found for intrinsic disorder^9,12^ or GC content^13^. Such results suggest the presence of an evolutionary filter that selects novelty with specific properties, that can only come from some parts of the genome and not others. More generally however, it is unclear exactly how the landscape of the noncoding genome shapes the landscape of emerging protein-coding novelty.

A recent study by Vakirlis et al.^14^ showed that, in the budding yeast *S. cerevisiae* genome the majority of intergenic regions carry the theoretical potential to, if translated, form transmembrane domains (TM domains). An increased tendency of thymine-rich regions, as are *S. cerevisiae* intergenes, to form TM domains is expected given that thymine-rich codons tend to code for hydrophobic amino-acids, stretches of which are predicted to form TM domains in protein sequences^15^. Yet the overall TM potential of *S. cerevisiae* intergenic regions is only partially explained by their nucleotide composition. Real intergenic sequences would, if translated, form significantly more TM domains than their randomly shuffled control counterparts, which maintain the same length and nucleotide composition.

This surprising finding ties in with the fact that *S. cerevisiae*-specific genes are enriched in TM domains, compared to conserved genes^6,14,16^. Furthermore, young genes that show beneficial fitness effects when overexpressed artificially are even more enriched in TM domains. These findings put together, led Vakirlis et al. to postulate a TM-first model for de novo gene birth, in which the cryptic potential to form TM domains is present in intergenic regions, and subsequently materializes during evolution. The unexpected discovery of TM-forming enrichment in intergenic regions of *S. cerevisiae* however was mostly left unexplained, as the focus of that study was elsewhere. Yet it raises some intriguing questions.

Is it simply a byproduct of underlying sequence properties, motifs or biases unrelated to coding sequences or does it actually have some protein-coding, evolutionary and biological role? Is this enrichment in TM-forming potential a general property of all budding yeast genomes? Is it evenly distributed on the genome, or does it come from specific regions/domains? Here, we took advantage of the availability of 332 budding yeast genomes annotated with a common pipeline, to perform a large-scale comparative analysis in an effort to answer these questions.

## Results

### Widespread enrichment in TM-forming potential in intergenic sequences of an entire subphylum

Given the unexpected enrichment in TM-forming potential found within the intergenic regions of *S. cerevisiae* (i.e. the fact that when computationally translated into proteins, they are predicted to form TM domains), we expanded the analysis to the genomes of 332 budding yeasts, representing the entire diversity of the subphylum *Saccharomycotina*^17^. For each genome, all intergenic regions were extracted and one artificial intergenic ORF (iORF) was generated from each one. TM domains were then predicted using Phobius and TMHMM for all iORFs longer than 6 nucleotides (see Methods for more details). In Figure 1A, we show, for each genome, the percentage of iORFs with at least one TM domain, as a function of the average intergenic thymine content (red points). As expected, there is a quasi-linear relationship. This translates to 25% - 75% of intergenes in most genomes having potential to form a TM domain. While this is partly expected, our analysis allows to fully appreciate the magnitude of this property.

**Figure 1.**
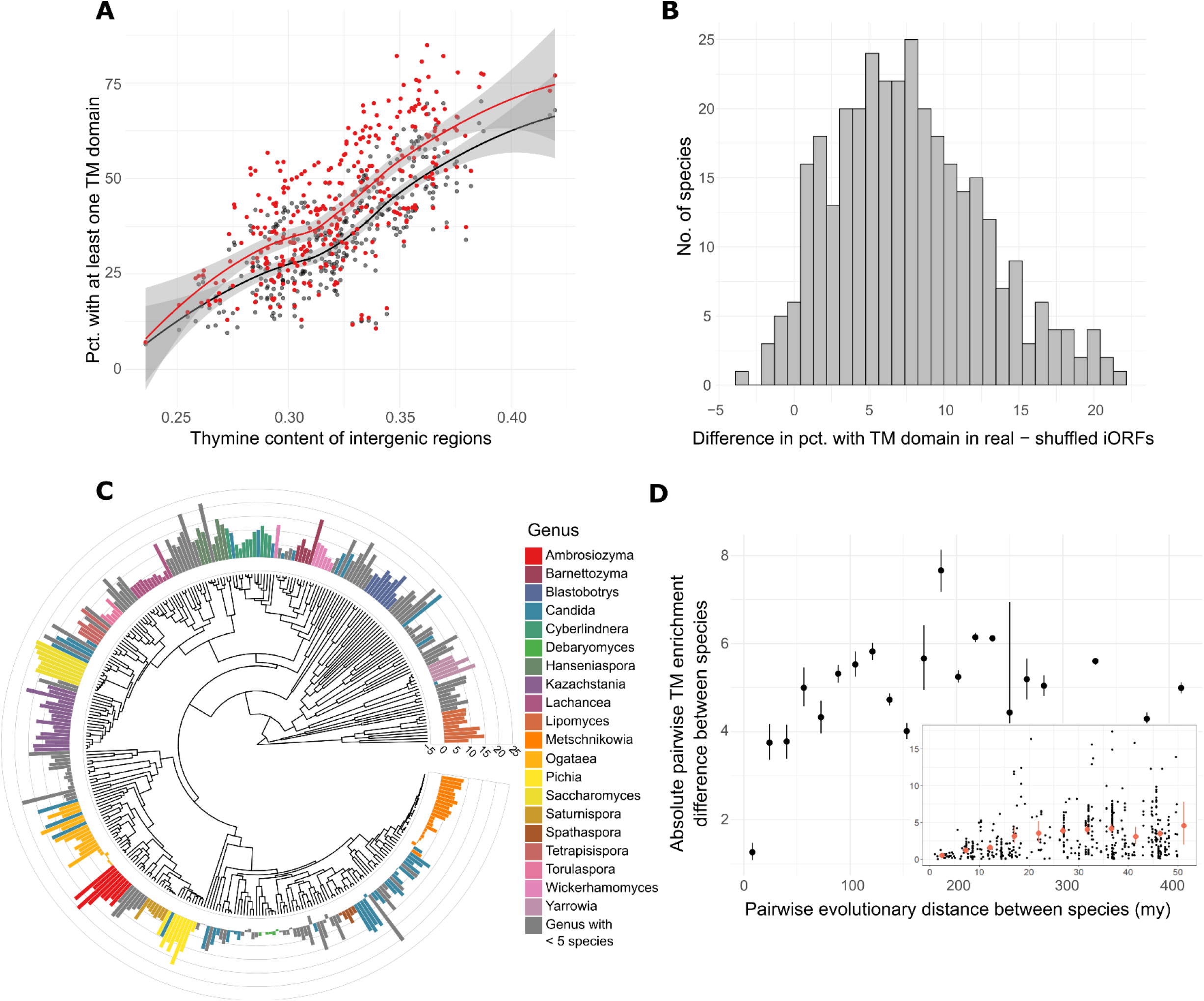
**A:** Genome-wide TM-forming potential of real (red) and shuffled (grey) iORFs as a function of the average thymine content of intergenes. Each species is represented by one red and one grey point. LOESS smoothing for each set is shown. **B:** Distribution of differences in percentage with TM domain between real and shuffled iORFs (red - black points in A). **C:** The same value as in B but plotted over the Saccharomycotina phylogeny, as reconstructed by Shen et al^17^. Genera comprising at least 5 species are colored. **D:** Distance in TM-forming enrichment values between pairs of species, as a function of the evolutionary distance separating the species (means with 95% confidence intervals in bins). Inset plot zooms in to the 0-50my region and shows all data points as well as means.

To understand whether thymine content may exclusively account for the predicted TM-forming potential, we generated randomized sequences with the same length and nucleotide composition of iORFs, as controls (hereafter shuffled iORFs, grey points in Figure 1A; see Methods). The real iORF distribution is overall higher than that of shuffled controls, and that is true for the entire range of thymine content. Figure 1B shows the distribution of the differences between these two percentages, using the Phobius predictions (TMHMM predictions strongly correlate with Phobius ones and can be found in Supp. Figure 1; Phobius will be used hereafter). There is substantial excess of TM-forming potential among real iORFs, with most genomes having between 3-12% more TM domains than expected by chance (as reflected in shuffled iORFs). In 309/332 cases the difference is positive and statistically significant (Supp. Figure 2A), while only one genome shows statistically significant depletion in TM-forming potential (*Kodamaea laetipori*). For brevity, we hereafter refer to this excess of TM-forming potential between real and shuffled iORF sequences, as TM-forming enrichment. Similar results were obtained when using the average percentage of sequence predicted to be TM (hereafter TM propensity) as a metric (Supp. Figure 2B).

We next asked whether the observed variation in this enrichment reflects the budding yeast taxonomy. In Figure 1C, we can see the distribution of TM-forming enrichment plotted over the Saccharomycotina phylogeny. We notice a clear taxonomic signal, with species from the same genus tending to have more similar values across species (ANOVA using genera as groups, P-value=1.981e-14; see also Supp. Figure 3). The variance within each genus does not correlate with the evolutionary distance separating the species (using any of the two different trees of Shen et al., P-value>0.28). Yet, species that are evolutionarily closer have more similar enrichment values (Figure 1D), even though the difference saturates quickly, at around 30-50my. This seems to suggest that, as species diverge during evolution, they also diverge, rapidly, in terms of this cryptic intergenic property. Interestingly, *S. cerevisiae* and the rest of the Saccharomyces genus represent the extreme end of the distribution (Supp. Figure 3). Thus, our analysis, which spans the diversity of all budding yeasts, reveals a much wider and heterogeneous picture, than the initial one of only a single species.

### TM-forming enrichment cannot be explained by compositional biases or presence of regulatory sequences

The observed enrichment in TM-forming potential could be a byproduct of the presence of promoters or other regulatory motifs within intergenic regions. Compositional biases or sequence patterns in regulatory regions could “accidentally” increase the odds to predict a TM domain, relative to a shuffled control. To investigate this possibility, we first tested whether removing regions likely to contain promoters had any effect in the observed difference. We generated a new dataset consisting of iORFs longer than 700nt (29.4% of all iORFs), from which we trimmed 200nt off each side (hereafter reiORFs). The TM-forming difference values of this reiORF set were highly correlated to the original one, with overall higher values (Figure 2A, Spearman’s Rho=0.79, P-value<2.2e-16; paired t-test P-value<2.2e-16 ; all iORF mean : 7.6, reiORF mean : 10.2), suggesting that promoter sequences are not responsible for the TM-forming enrichment. In fact, the 200nt edges show significantly weaker TM-forming enrichment, even compared to similarly sized iORFs generated from entire intergenes (Supp. Figure 4, left panel). As an additional control, we tested an even stricter version of reiORFs, taking only iORFs >1500nt in length (10% of the total) and trimming 500nt off each side, and obtained similar results (Supp. Figure 4, right panel). If anything, then, the enrichment seems to come more from the middle regions of intergenes than from sequences likely to have regulatory roles.

**Figure 2.**
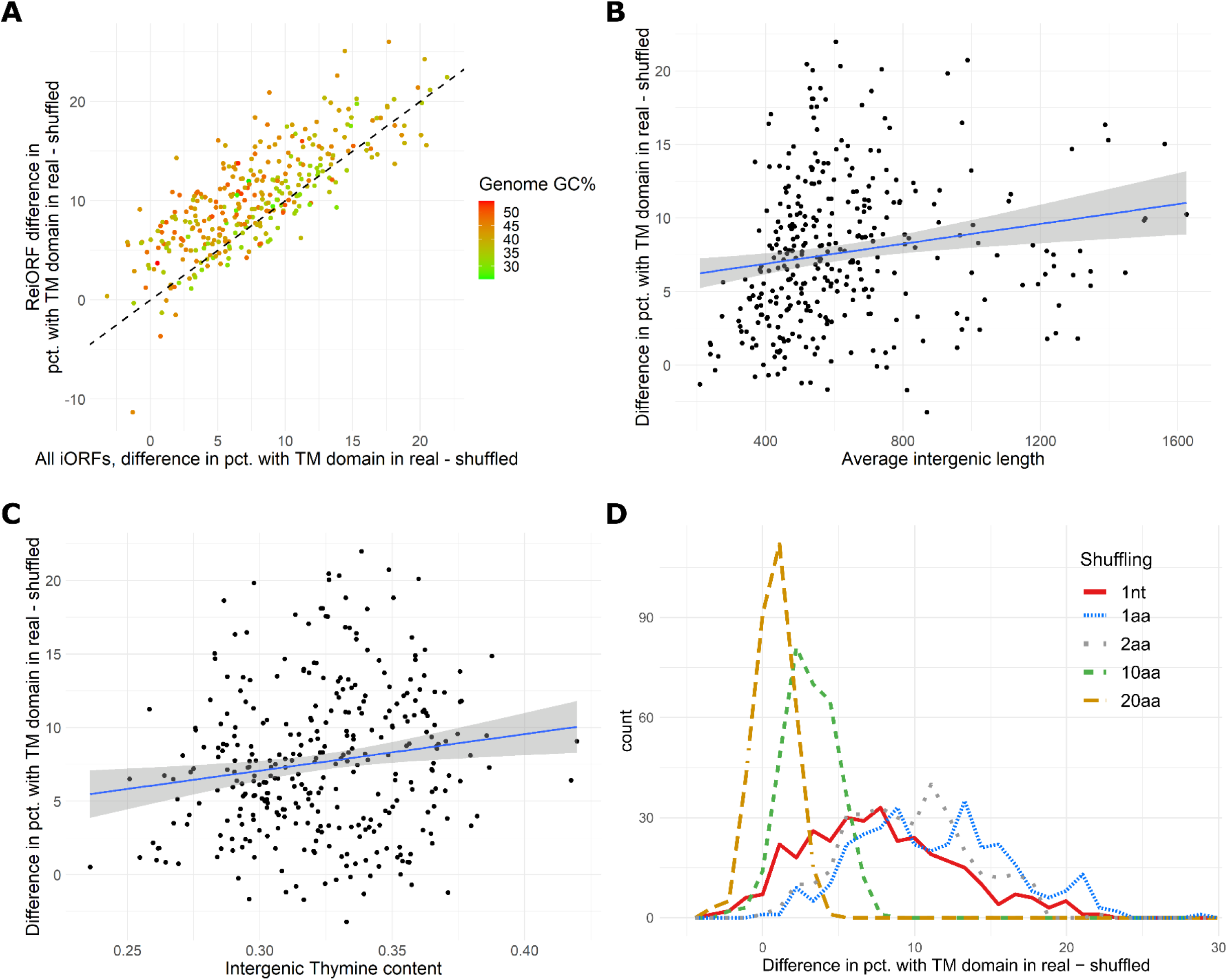
**A:** Correlation between TM-forming enrichment in all iORFs and reiORFs (iORFs longer than 700nt with 200nt trimmed off each edge). Points are colored by genomic GC%, showing that the correlation holds over varying GC% values. **B:** TM-forming enrichment as a function of average intergenic Thymine content. **C**: TM-forming enrichment as a function of average iORF length. **D:** Distributions of TM-forming enrichment as measured based on single nucleotide shuffling (default), 1aa, 2aa, 10aa and 20aa shuffling.

Given that the TM propensity of a sequence is largely driven by its thymine content, we next asked whether TM enrichment could be explained by differences in the nucleotide composition of genomes. We found only a weak correlation to the average thymine content (Rho=0.16, P-value=0.004, Figure 2B) or GC content (Rho=-0.18, P-value=0.00093) of intergenic regions, or to the average GC content of entire genomes (Rho=-0.2, P=0.0002). Indeed, as previously mentioned, even though more real iORFs have TM domains, the TM-forming potential in both real and shuffled iORFs is correlated similarly to thymine content across budding yeasts (Figure 1A). A moderate correlation was found to the average length of intergenic regions (Rho=0.27, P-value=7.9e-7; Figure 2C), but this seems to be an artefact due to the paucity of TMs predicted in very small intergenic sequences, which we have included (the vast majority of TM helices are longer than 10aa^18^), and disappears when we exclude them (P-value=0.95; Supp. Figure 5A). Weaker correlation was found to the alternative metric, that is the difference in the average percentage of iORF sequence predicted to be TM (Rho=0.19, P-value=0.0004; Supp. Figure 5B).

An alternative possibility is that this signal is coming from motifs of more than one nucleotide which are split apart when the sequences are being shuffled. To test whether this is true, we repeated our shuffling using one codon and two codons (i.e. 1aa and 2aa after translation) as the unit being shuffled, instead of single nucleotides, and obtained similar results (Figure 2D). In fact, both alternative sequence permutations resulted in even deeper enrichment, that is, even lower TM-forming potential in the shuffled sequences compared to the real ones (mean difference in pct. with TM domain: 7.65%, 11.4%, 9.6% for 1nt, 1aa and 2aa shuffling respectively). In contrast, and as expected given the average size of TM domains, increasing the shuffling unit to 10aa dramatically reduces TM-enrichment (avg. 3.1%), while increasing to 20aa effectively eliminates it (avg. 0.75%; Figure 2D). This confirms that sequence motifs that underlie this TM-forming potential indeed approximate the length of entire TM domains.

Along the same lines, a major determinant of the TM propensity of a protein sequence is its hydrophobicity. It follows that the signal we detect at the TM domain level, could be reduced to the hydrophobic content of proteins (more TM domains in real iORFs would simply be due to more hydrophobic amino-acids in real iORFs). This is however not the case, as the difference in average hydrophobicity (real – shuffled iORFs) shows a negative correlation to the difference in percentage with TM domains (Rho=-0.31, P-value=5.2e-9). Strikingly, this means that real iORFs have at the same time both slightly lower hydrophobic potential (paired t-test P-value<2.2e-16; Cohen’s d = -1.28) and higher TM-forming potential (Figure 3A). In other words, they make more with less (using alternative scales to calculate hydrophobic content results in similar differences, see Supp. Figure 6). Note that the difference in hydrophobicity between real and shuffled iORFs, correlates negatively to the thymine content of iORFs (Figure 3B; Rho=-0.57, P<2.2e-16). This means that in genomes with higher thymine content, shuffling of iORFs will actually increase their hydrophobic content. This is in stark contrast to TM domain difference, where only a weak, positive correlation was found to thymine content (Figure 2B). Thus, hydrophobicity as a property does not explain the observed TM-forming enrichment, while at the same time showing a consistent, more predictable trend.

**Figure 3.**
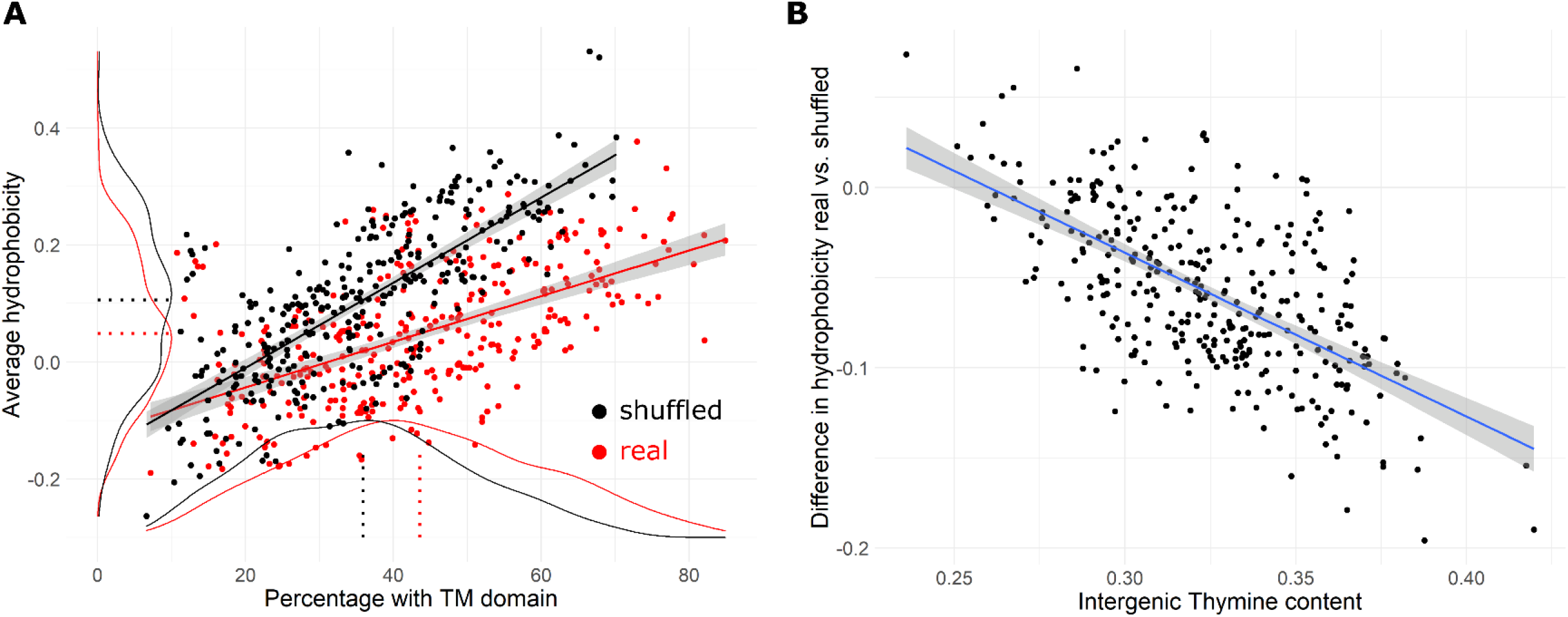
**A:** Percentage with at least one TM domain in real and shuffled iORFs plotted against their average hydrophobicity. Dotted lines show distribution means. Density plots have been scaled for visual purposes. **B:** Difference in hydrophobicity between real and shuffled iORFs is strongly negatively correlated to the average intergenic thymine content.

### Investigation of TM-forming enrichment within individual genomes

So far, we have focused on average values of entire genomes. Zooming in to the individual iORF level and to their distribution within genomes could help us trace the enrichment to specific genomic regions. We analyzed six genomes selected to represent extremes in TM-forming enrichment, GC% and coming from different genera (Supp. Table 1). Examining how the difference in TM propensity between real and shuffled iORFs is distributed across the genome revealed no obvious hotspots and no distinct positional bias (Figure 4A and B for *S. uvarum*, see Supplementary Figure 7 and 8 for the other 5 species). No strong correlation to either individual iORF thymine content or length was found in any genome we investigated (all Pearson’s R correlation coefficients < 0.11). In contrast to what we found when comparing genome-wide averages, there is weak, but statistically robust correlation of difference in TM-forming propensity to difference in hydrophobicity (Figure 4C and Supp. Table 1). This means that at the intra-genomic level we do see that an enrichment in hydrophobicity underlies the TM-forming enrichment, but it can only explain a minor part of the variation overall.

**Figure 4.**
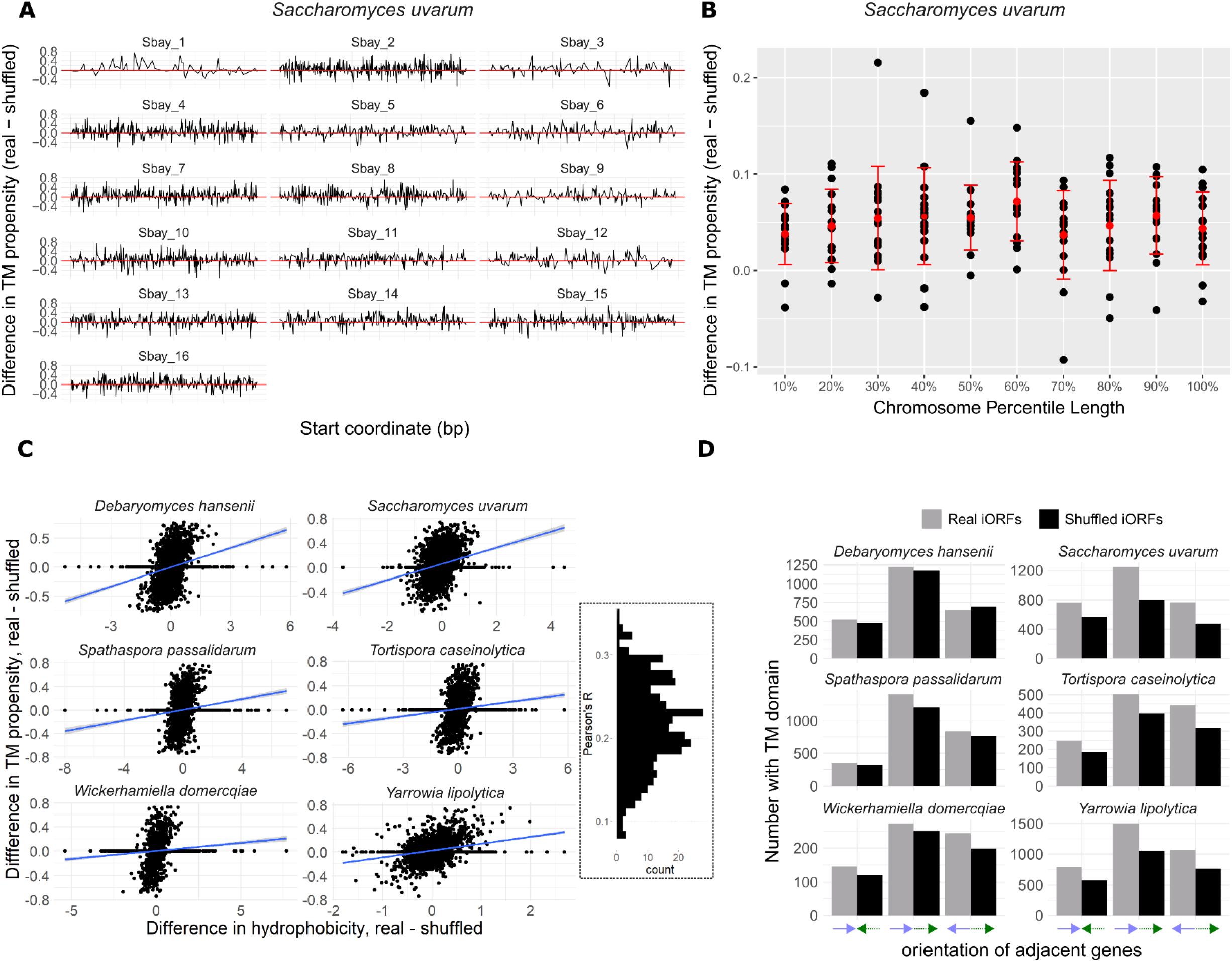
**A:** Distribution across the genome of difference in TM-propensity (percentage of sequence predicted to be TM) for *S. bayanus*, which has one of the highest TM-forming enrichment values (20% more real iORFs have TM domains than shuffled ones). No hotspots or patterns can be observed. **B**: Same value as in A, but each chromosome is broken into 10 bins of equal length and an average value is calculated for each one. No strong positional bias can be seen (means and confidence intervals shown in red). **C**: Only a weak correlation can be found between difference in hydrophobicity and difference in TM propensity, in the six species we investigated. Over all 332 species, correlation coefficients range from 0.08 to 0.35 (histogram in dotted rectangle). **D**: No significant effect of adjacent gene orientation on TM-forming enrichment between real and shuffled iORFs (difference between grey and black bars).

We next reasoned that, since we can predict the enrichment of an intergenic region in regulatory sequences based on the orientation of the adjacent genes, iORFs coming from regions flanked by differently oriented genes might show different trends (e.g. an intergene flanked by two genes in opposing orientation should in theory contain no promoter sequences). Yet TM-forming enrichment is similar in all 3 arrangements (Figure 4D). Overall, these findings strongly suggest that this property encompasses the entire genome in a relatively uniform manner. Finally, using the data from Vakirlis et al. in *S. cerevisiae*, we checked whether TM-rich iORFs are situated next to annotated proteins with high TM content but found no meaningful association (Supp. Figure 9).

### Species with higher TM-forming enrichment in intergenic regions tend to have more TM-rich young genes relative to conserved ones

In *S. cerevisiae* it was previously shown that young genes are significantly TM-richer than conserved ones. This difference, taken together with the even higher TM-forming potential of intergenic regions, led to the proposal of a gradual drop of TM propensity as novel genes emerge out of intergenic regions and mature into conserved ones. But whether this is a general trend, or perhaps even a rule that might hold across different species, is still unknown.

To provide an answer, we predicted TM domains in a set of Saccharomycotina proteins that originated less than 30mya, based on the most recent common ancestor of the species in which they were found (see Methods). We did the same in a subset or proteins conserved inside and outside of the subphylum, and thus correspond to ancient proteins, which we subsampled to have similar length distribution as young ones (see Methods). We found that in most yeast species, we see the opposite trend than in *S. cerevisiae*: young genes are less TM-rich than conserved ones, albeit with significant variability even among closely related species (Figure 5A). *S. cerevisiae* is once again an outlier, exhibiting the second-highest difference in TM propensity in young genes compared to conserved ones.

**Figure 5.**
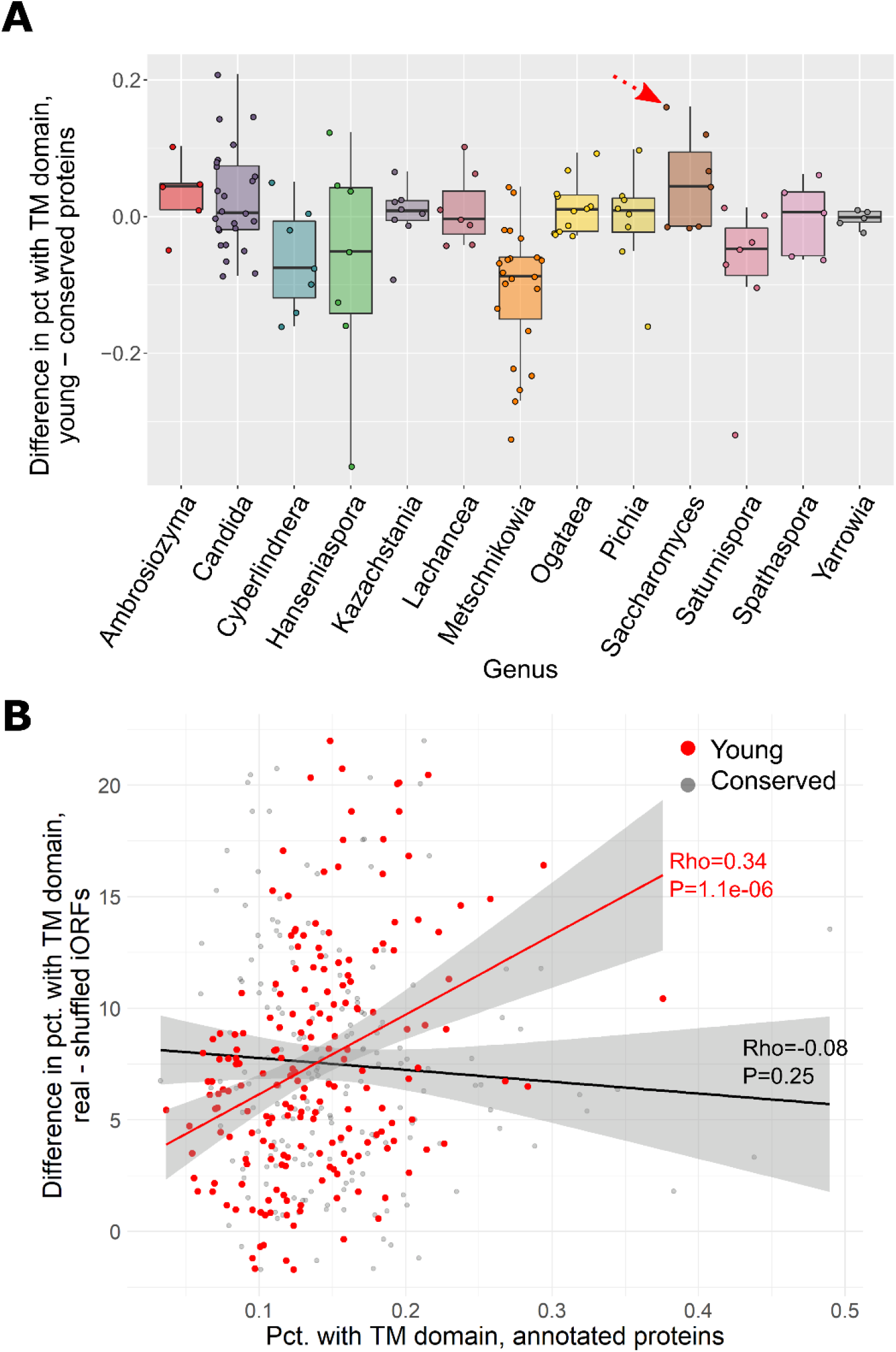
**A:** Difference in percentage with TM domain between young and conserved genes across genera, showing that, in that aspect too, *S. cerevisiae* is an outlier (red arrow). Conserved genes have been subsampled to match the length distribution of young genes. Only species with at least 20 young genes are included in this analysis. **B:** The TM-forming enrichment of iORFs correlates to the TM propensity of young genes, but not of ancient conserved genes, across species (represented by one grey and one red point each).

With this observation at hand, we next looked for possible links between the excess TM potential of intergenic regions and the TM richness of young genes. Is that excess potential translated into more TM rich genes through the emergence and evolution of novel genes? By plotting the TM-forming enrichment of iORFs in each species to the average TM propensity in its young genes, we see a clear increasing trend (Spearman’s Rho=0.34, P-value=1.1e-6; Figure 5B, red points). This trend is absent in conserved genes (P=0.25) and also significantly reduced if we replace TM-forming enrichment with the TM-forming potential of real iORFs (Rho=0.17, P-value=0.02). In other words, it is the enrichment relative to the theoretical expectation that exhibits the stronger correlation, and not the actual TM content of iORFs. This seems to suggest that in species where this enrichment is stronger, it somehow gets translated into young genes that tend to encode more TM domains.

## Discussion

The noncoding genomic landscape is expected to influence the process of emergence of novel genes, and understanding the different aspects of this influence is crucial. Drawing motivation from the recent discovery of widespread cryptic potential to form TM domains, found in the genome of *S. cerevisiae*, we performed an expanded, comparative investigation of this property in hundreds of available budding yeast genomes. The relevance to the process of de novo gene emergence is obvious, especially given the initial findings from *S. cerevisiae*. Yet, zooming out to the subphylum level revealed a complex and heterogeneous picture. Additional work is now necessary to substantiate the nature of the relation of TM-forming enrichment to the birth of new genes.

Previous analyses have shown that other basic structural units of proteins beyond TM domains could readily be produced from random and intergenic sequences^19^. This was best shown recently by Papadopoulos et al.^5^, who analyzed small intergenic ORFs and showed that they display a wide diversity of potential protein folds, which are also found in real proteins. Our work here, focused on the enrichment relative to shuffled controls, opens the door to similar analyses of other structural units, such as beta sheets or intrinsically disordered regions. This could potentially reveal some underlying organization, related to a previously unappreciated aspect of the biology of genomes.

We should nonetheless also keep in mind that this enrichment might turn out to have no biological significance. For now, this is only based on computational prediction of artificial protein sequences. Before we can confidently claim anything, a more in depth, direct analysis of the link of TM-forming enrichment to the process of de novo gene emergence is necessary, which would combine comparative genomics, transcriptomics and ribosome-profiling techniques. Additionally, the TM-forming potential and enrichment of iORFs should be confirmed experimentally and at a large-scale, to corroborate the results of the computational predictions.

Despite our efforts, we were not able to find a property that strongly correlates to TM-forming enrichment, and which would help explain its origins. We established that the observed signal does not come from GC content, promoter regions or simply the presence of potential hydrophobic amino acids. This may be suggestive of an inherent sequence structure that underlies the iORFs propensity to form TM domains, which cannot be directly attributed to either nucleotide or amino-acid content. Existing intergenic sequences appear to be somehow tuned to allow for the encoding of TM domains, even with thymine and hydrophobic residue contents that are smaller than expected. It is thus due to something else, some underlying cause that leads to stretches of hydrophobic amino acids, within existing, seemingly unconstrained, intergenic regions.

On the one hand, it is possible (and more easily conceivable) that some feature or property of intergenic regions that has escaped us so far, and is unrelated to protein-coding potential, is responsible. This would mean that any influence this property has on potential novel genes has come about indirectly, as a byproduct of signals with an entirely distinct evolutionary purpose (e.g. regulatory).

On the other hand, we might speculate that the source of this enrichment has no other evolutionary purpose but functional predisposition of novelty. It is conceivable that weak selection forces could have acted to instill optimal levels of TM forming potential within intergenic regions. This could boost the ability of genomes to produce novelty and potentially make organisms more robust, and more evolvable. For now, this can only be purely speculative. Similarly, this enrichment could reflect evolutionary pressure to avoid expression of unwanted products, balanced against the advantage of occasionally allowing novelty to emerge. The overrepresentation of TM domains, perhaps coupled with the absence of other signals, could be part of a surveillance mechanism that detects and degrades aberrant translation products, similarly to what was found in a recent preprint^20^.

A third explanation could be that this enrichment has its source in the past, rather than being, in a sense, evolutionarily future-looking. One can imagine that remnants of ancient TM proteins (or domains or existing ones), which are known to evolve fast and be easily lost^21^, could ultimately leave behind the sequence motifs that we detect. However, since we saw that the enrichment is distributed across the genome, such remnants would have to become scattered across the genome over time, making this scenario more complicated. Furthermore, we were unable to find evidence of co-localization of TM-forming potential in iORFs to TM containing proteins in the *S. cerevisiae* genome. Nonetheless, further work is now needed to conclusively refute or confirm this hypothesis, and more generally to elucidate the origins of the TM-forming enrichment.

## Materials and Methods

### Data collection

332 Saccharomycotina yeast genome assemblies, annotations, CDS and protein sequences were downloaded from the repository provided by Shen et al^17^. From the same repository, we downloaded the two available Saccharomycotina phylogenies, as well as homologous protein families.

### Generation of real and shuffled iORFs

Artificial intergenic ORFs (iORFs) were generated out of every intergenic region larger than 6 nucleotides as in ^13,14^. For every genome assembly, coordinates of intergenic regions were obtained using the BEDtools^22^ *subtract* tool together with the corresponding annotation GTF file. We extracted the intergenic sequences, removed stop codons in the +1 reading frame and then translated the sequences using custom Python scripts (one intergenic sequence produced one iORF). Sequences that had non-ACGT nucleotides were discarded.

To construct the initial, single nucleotide shuffled iORF controls, nucleotide iORF sequences were shuffled using custom Python scripts, following the stop codon removal. Each position of the nucleotide sequence was randomized and whenever an in-frame stop codon was formed from the randomization, its 3 positions were randomized again until they did not form a stop codon. This ensured that the shuffled sequences retained the exact same nucleotide composition as their real counterparts. For 1,2,10 and 20 amino-acid shuffled controls, sequences were manipulated accordingly, and no additional stop codon removal was necessary.

A separate dataset of iORFs (reiORFs) was generated by taking all intergenic regions longer than 700 nucleotides and trimming 200 nucleotides from each edge to retain a sequence consisting of at least 300 nucleotides in the middle. Shuffling and translation was then applied as detailed above. The same was applied in order to generate an additional subset out of iORFs longer than 1500 nucleotides, where 500 nucleotides from each edge were trimmed.

### Prediction of protein and CDS properties and various statistics

Transmembrane domains were predicted locally with Phobius^23^ and TMHMM^24^. Grand Average of Hydrophobicity index (GRAVY) on the Kyte-Doolittle scale was calculated using *codonw*^25^ and on two additional scales (Miyazawa and Wolfenden) using the R package *peptides*^26^. Length, thymine content and GC% of iORFs were calculated using in house Python scripts. Based on the homologous protein families of the Saccharomycotina reconstructed by Shen et al., we calculated ages of origin for each one, by determining the branch of the most recent common ancestor of all the species in which the family was present. We defined young families as those that originated at most 30mya. To build the conserved protein set, we first randomly selected 1000 families, present in almost all Saccharomycotina species and with members in outgroups outside of Saccharomycotina. To be able to compare the young and conserved set, we sampled the conserved set to obtain a subset with length distribution comparable to the young one. Subsampling was done separately for each species, and only for species for which at least 20 young proteins existed. Subsampling was performed with replacement, using a version of inverse transform sampling, as previously described^14^. An alternative method that minimizes sampling the same genes multiple times was also applied and produced identical results (data not shown). All statistical comparisons were performed in R v. 3.6.2. Plots were generated using *ggplot2*^27^ and *ggtreeExtra*^28^.

## Acknowledgements

We thank the members of the Computational Genomics Group for fruitful discussions which contributed to the development of this work.

## Author information

### Author contributions

NV and CN conceived the study. NV and CN supervised the work. NV and ET performed the analyses. NV, CN and ET analysed the data. NV, CN and ET wrote the paper. All authors read, finalized and approved the final manuscript.

### Corresponding authors

Correspondence to Nikolaos Vakirlis and Christoforos Nikolaou

## Supplementary Figures

**Supplementary Figure 1 :**
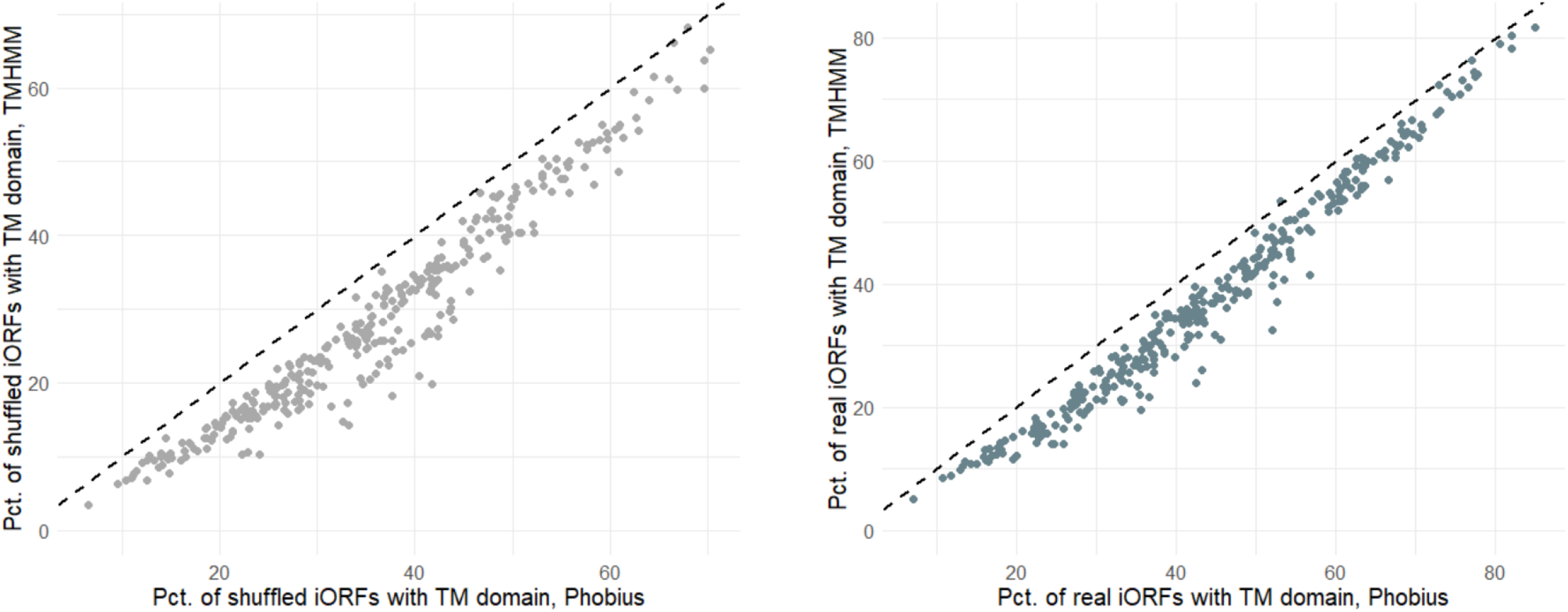
Correlation between percentages with TM domain in each species as predicted by Phobius and TMHMM, in shuffled (left) and real (right) iORFs. Values are highly correlated with Spearman’s Rho of 0.97 and 0.99 respectively.

**Supplementary Figure 2:**
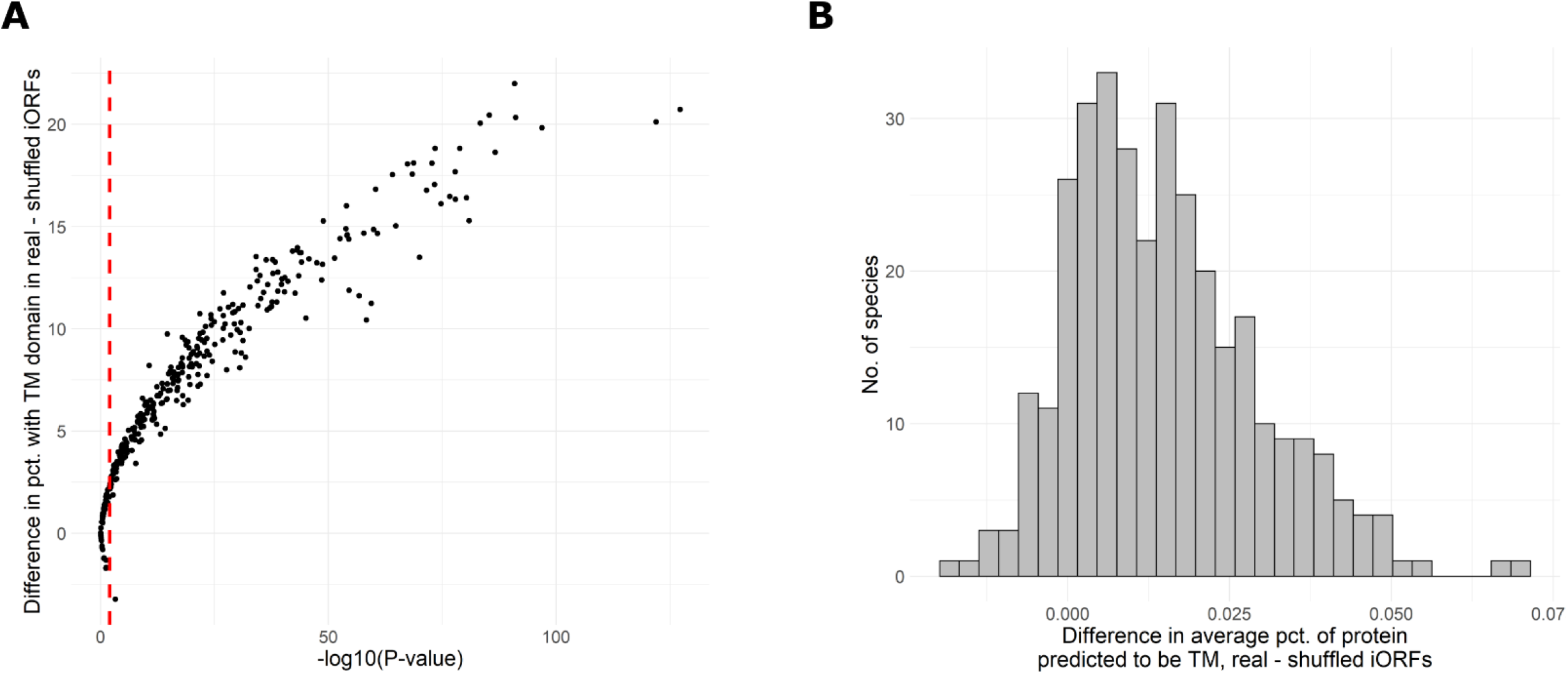
A: Fischer test P-values for comparisons of percentages with at least one TM domain between real and shuffled iORFs. B: Distribution of values of difference average percentage of iORF protein predicted to be TM, between real and shuffled iORFs of each species.

**Supplementary Figure 3:**
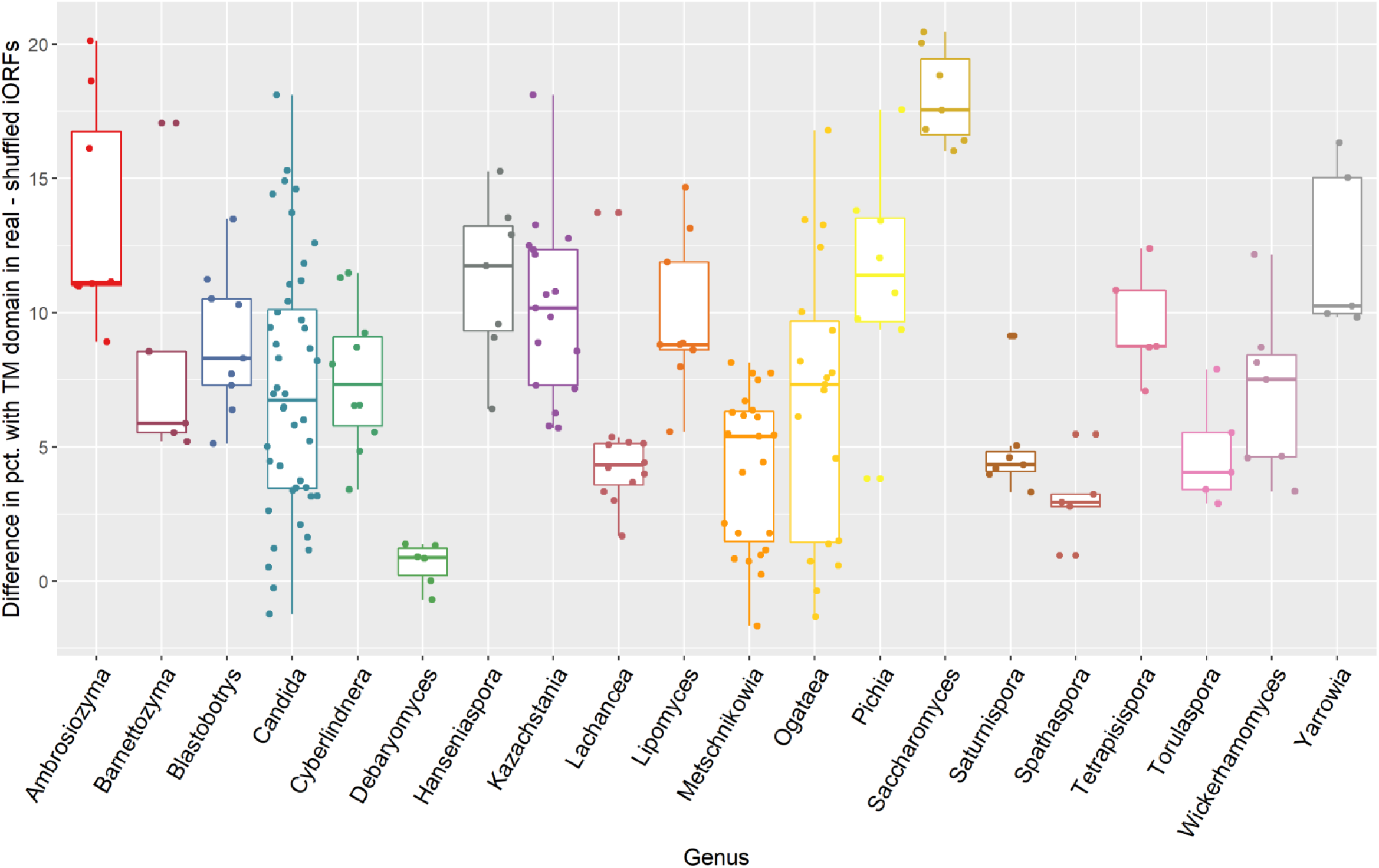
Distribution of difference in percentage with TM domain between real and shuffled iORFs (TM-forming enrichment) across various Saccharomycotina genera. Only genera with at least 5 species are shown. The Saccharomyces genus has by far the highest median of all genera.

**Supplementary Figure 4.**
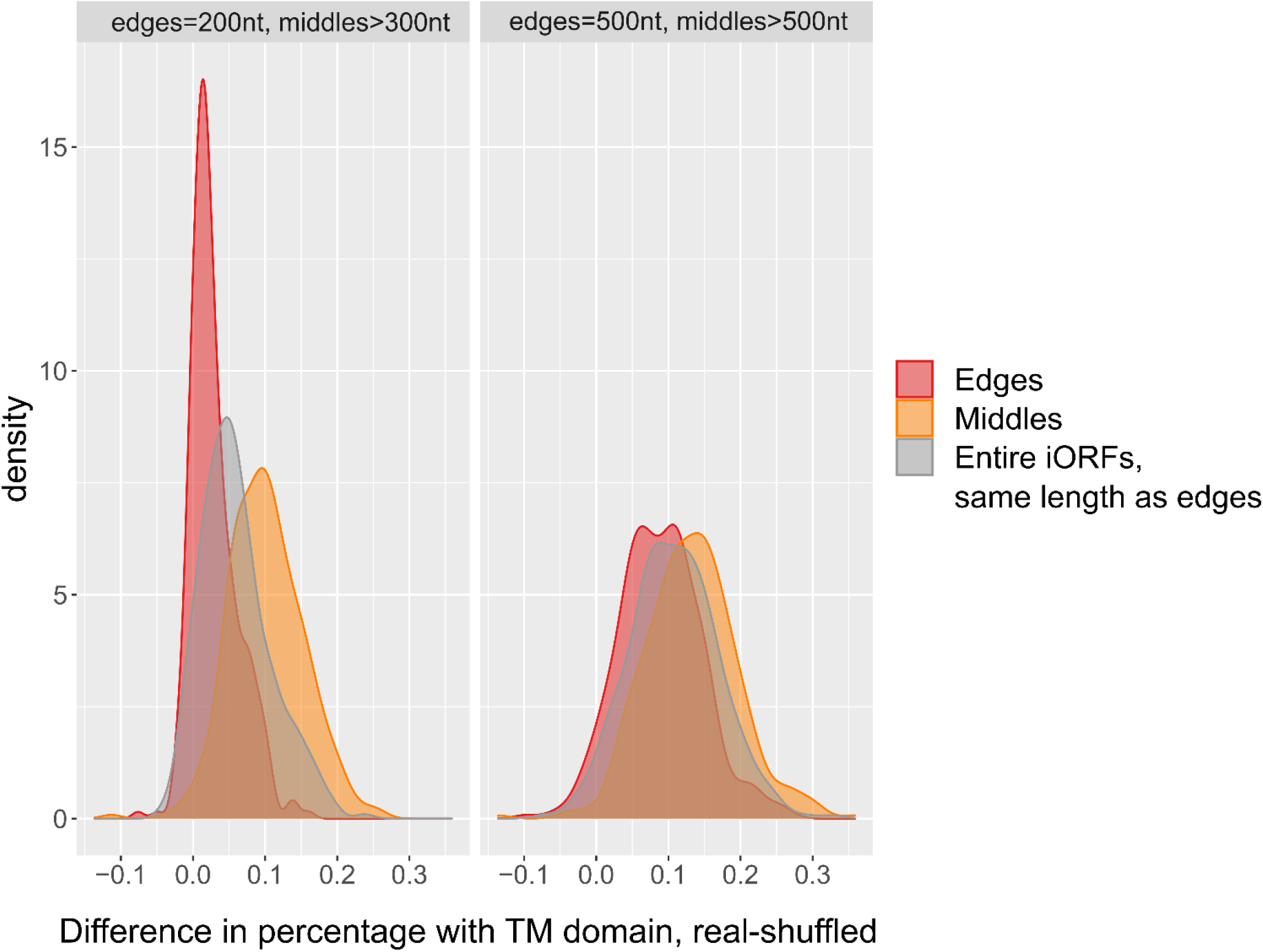
Density plots of TM-forming enrichment found in edges and middle parts of two versions of reiORFs, as well as entire iORF controls with similar length average as edges.

**Supplementary Figure 5.**
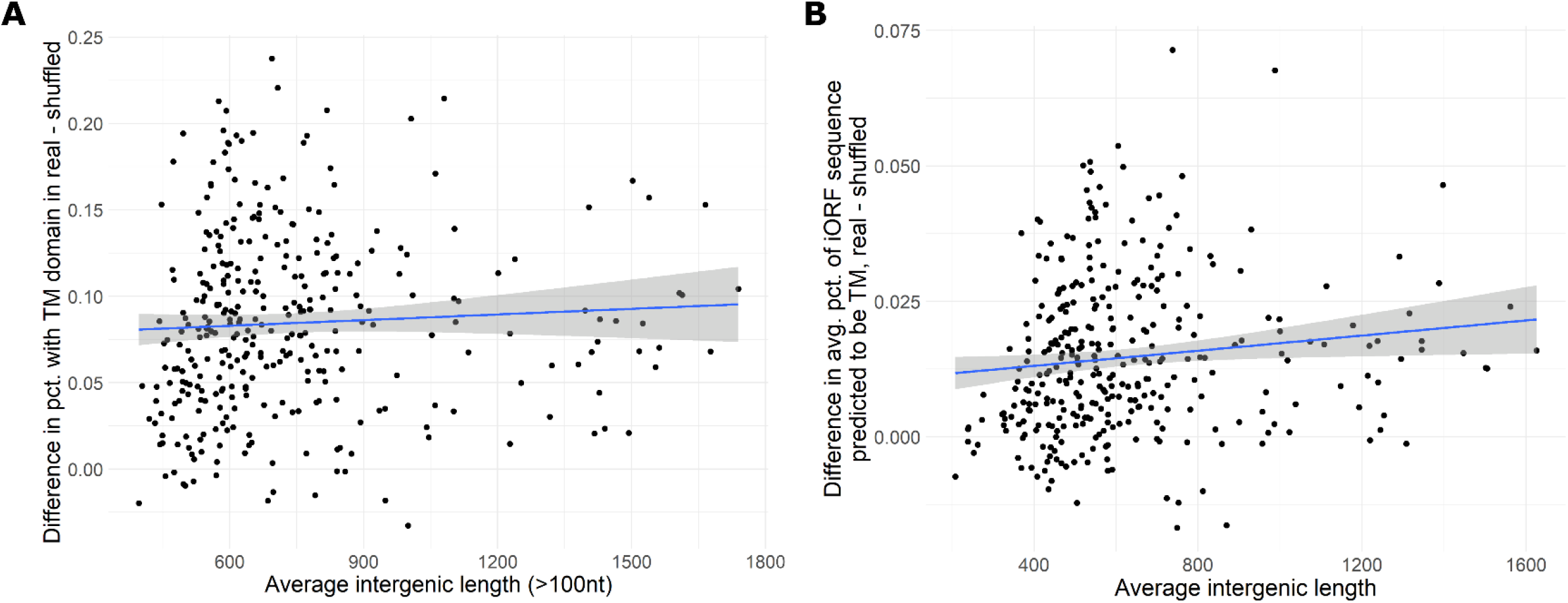
A: No correlation between TM-forming enrichment and average iORF length when excluding intergenes shorter than 100nt, for which TM formation is less likely. B: A weak correlation is found when correlating the difference in the average percentage of sequence predicted to be TM to the average iORF length, even over the entire dataset (Rho=0.19, P-value=0.0004).

**Supplementary Figure 6.**
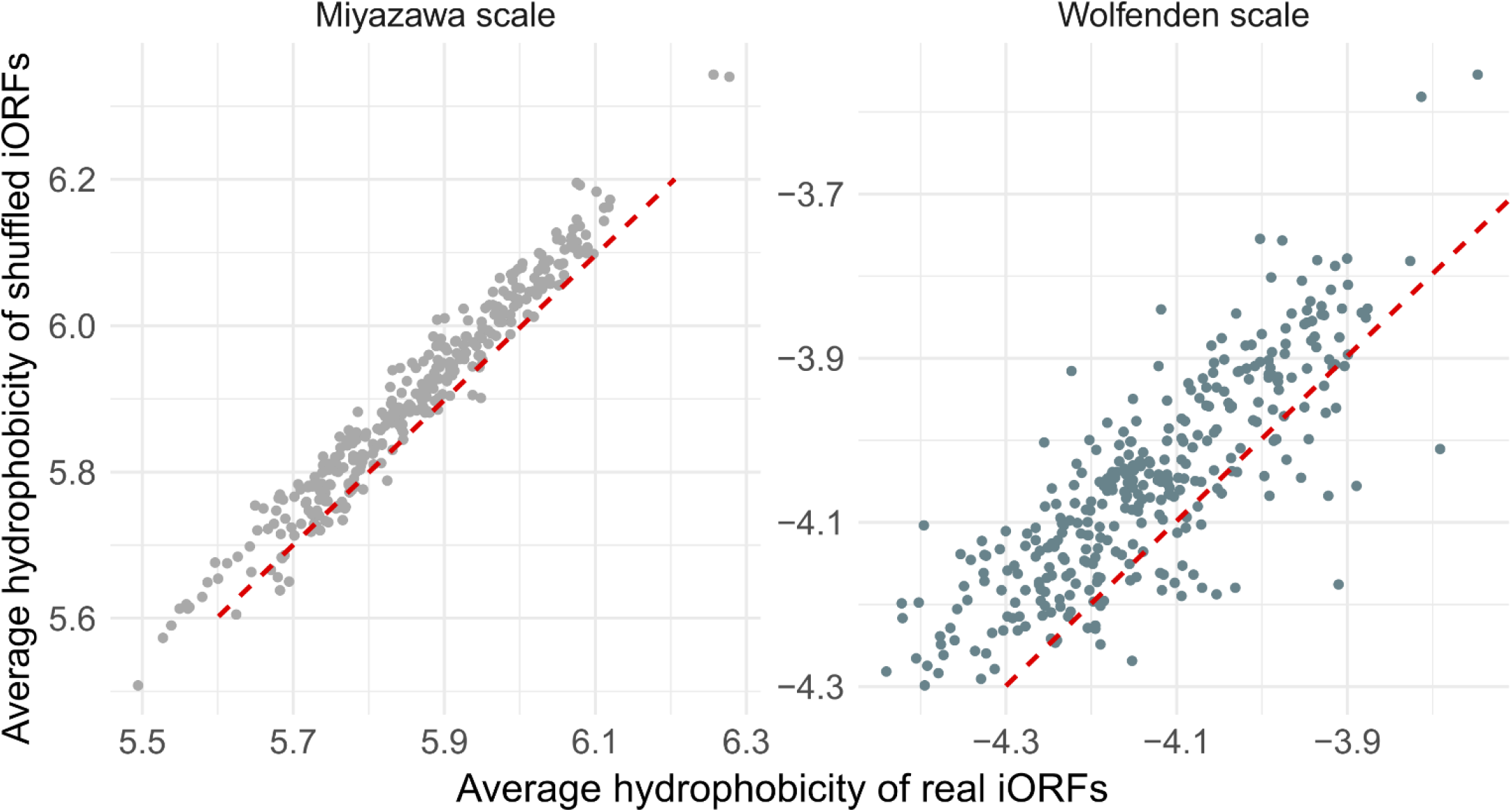
Hydrophobicity of real and shuffled iORFs as calculated by two alternative scales. Red line shows 1-to-1 relationship (slope=1, intercept=0). Paired t-test P<2.2e-16 for both; Cohen’s d -1.3 and -1 respectively.

**Supplementary Figure 7.**
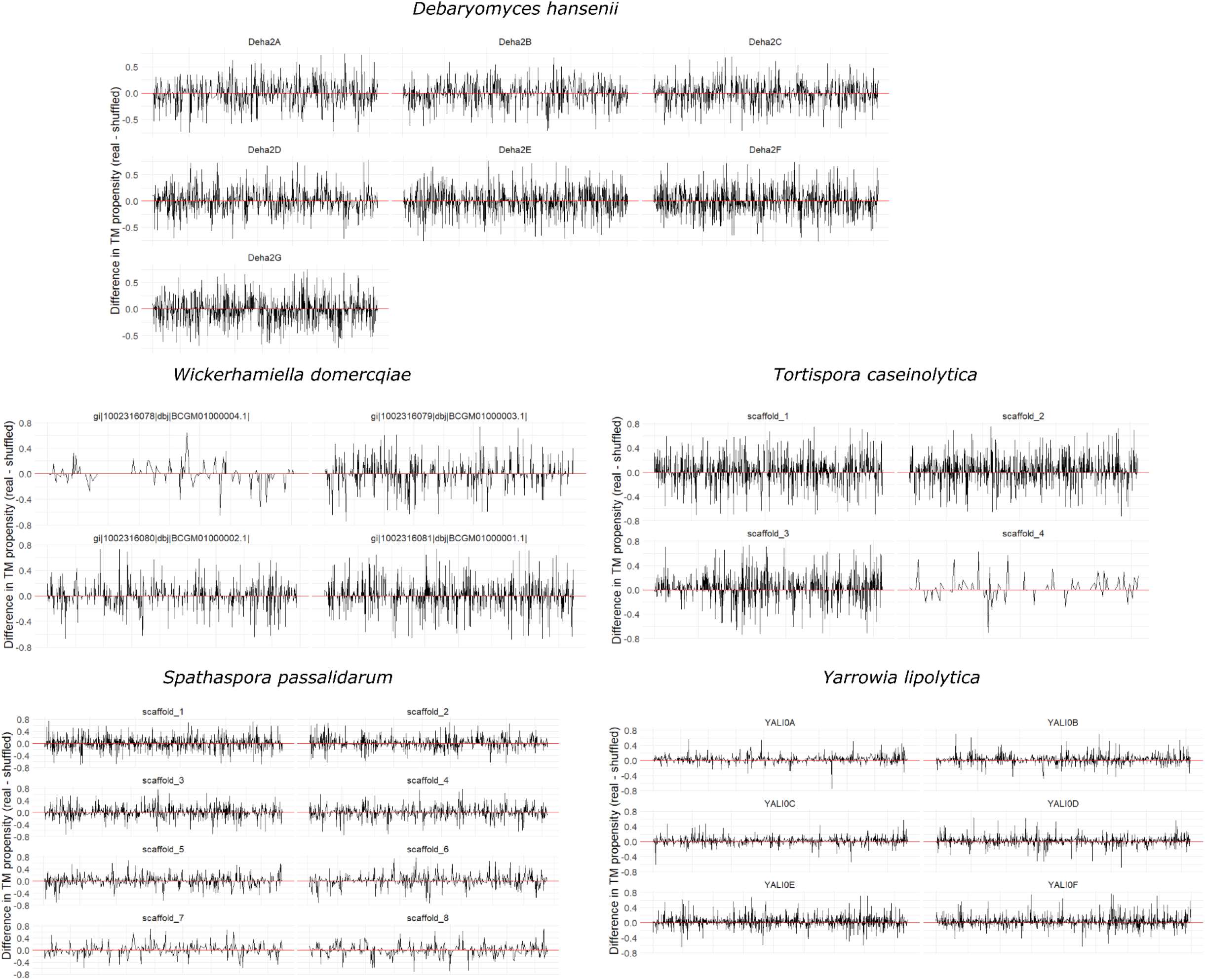
Difference in TM propensity of individual iORFs (percentage of protein predicted to be TM in real - shuffled sequences) across the genomes of 5 Saccharomycotina species.

**Supplementary Figure 8.**
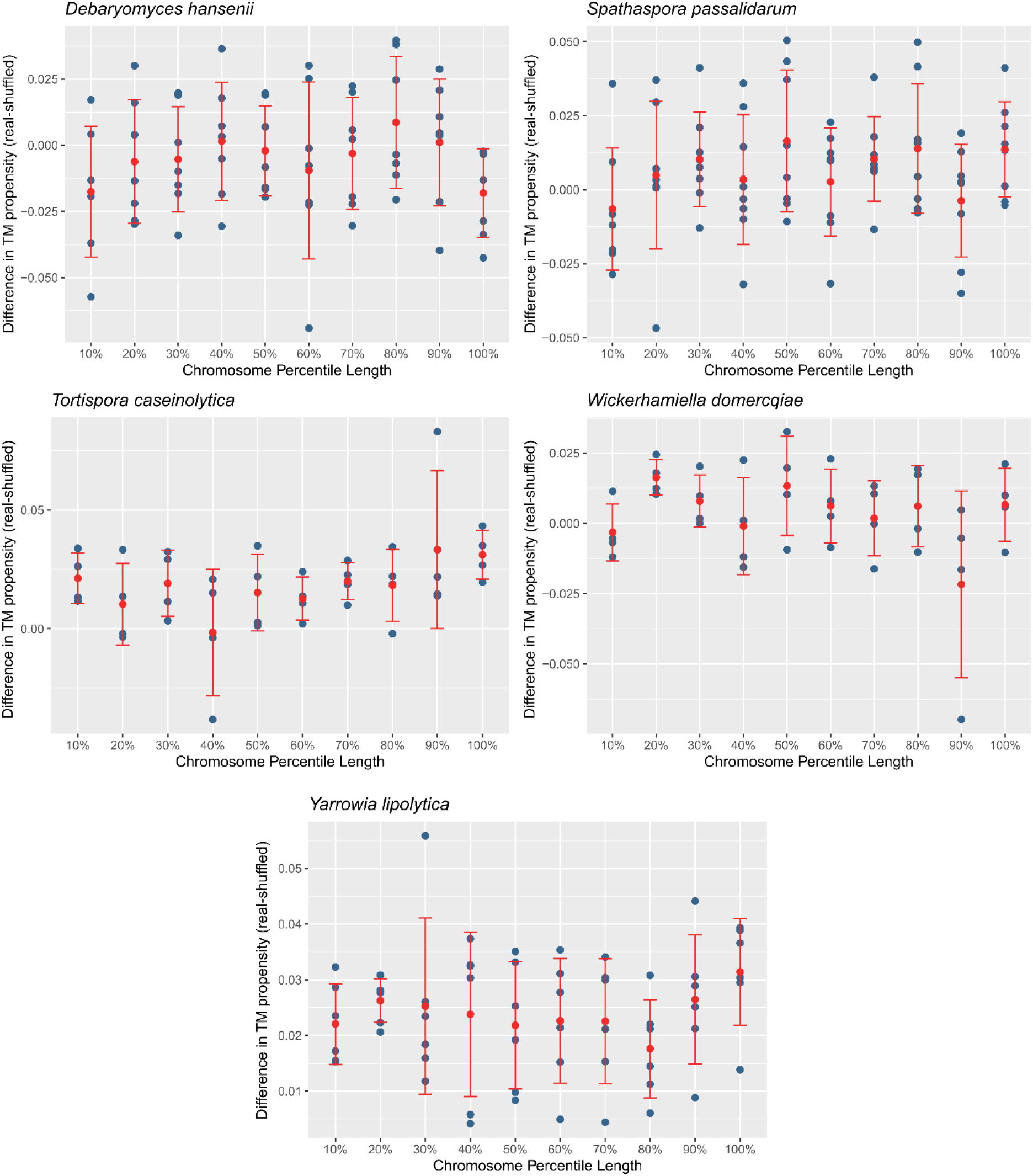
Distribution of difference in TM propensity (percentage of protein predicted to be TM) between real and shuffled iORFs, in chromosome percentile length bins.

**Supplementary Figure 9.**
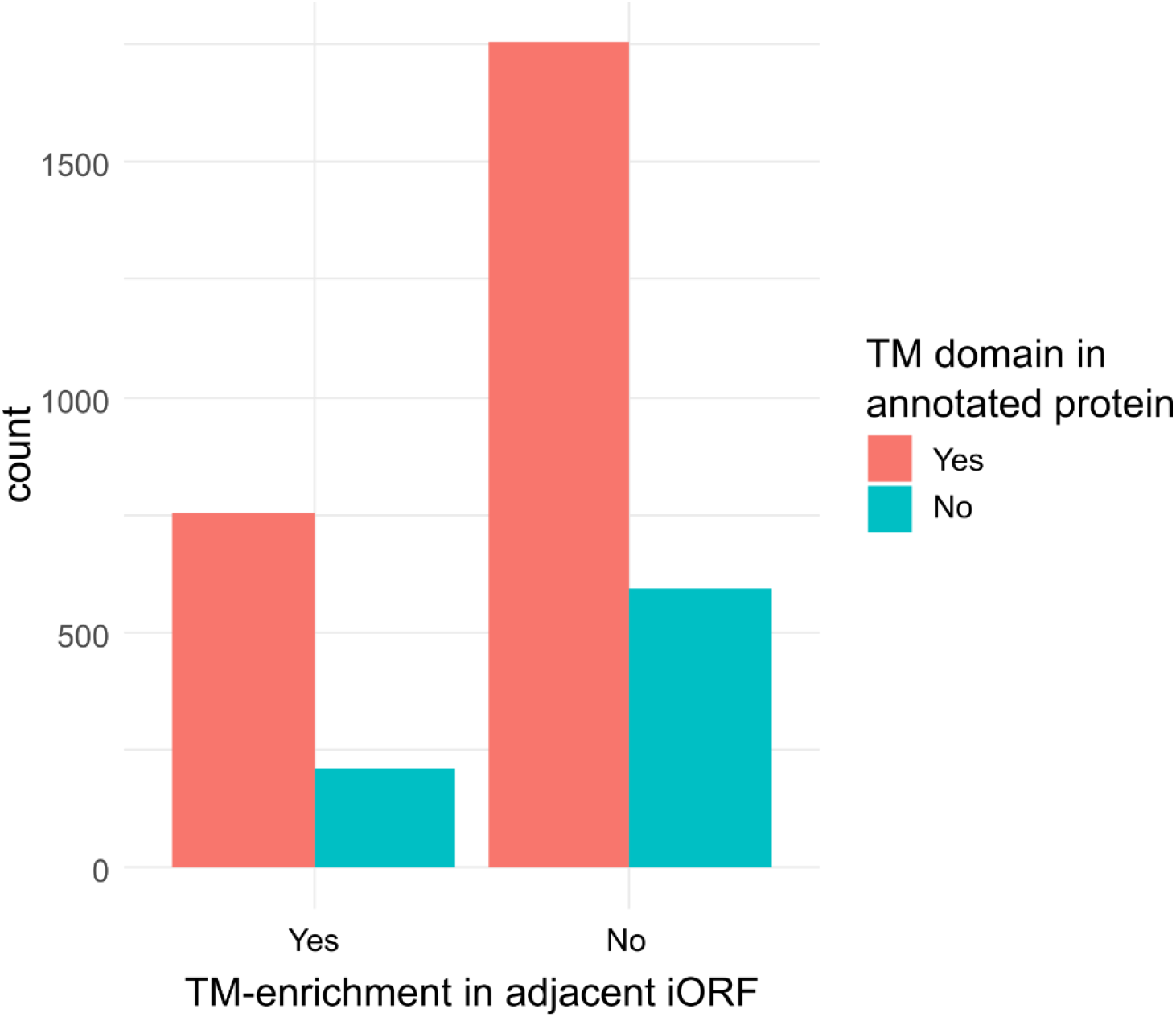
Association between TM-enrichment of iORFs and presence of TM domains in adjacent annotated proteins in *S. cerevisiae*. A negligible effect can be found that is marginally statistically significant (X^2^ P-value = 0.04).

**Supplementary Table 1**: Genomic information on the 6 Saccharomycotina species analyzed at the intragenomic level.

